# Rare variants in dynein heavy chain genes in two individuals with *situs inversus* and developmental dyslexia

**DOI:** 10.1101/2020.03.30.011783

**Authors:** Andrea Bieder, Elisabet Einarsdottir, Hans Matsson, Harriet E. Nilsson, Jesper Eisfeldt, Anca Dragomir, Martin Paucar, Tobias Granberg, Tie-Qiang Li, Anna Lindstrand, Juha Kere, Isabel Tapia-Páez

## Abstract

**Background:** Developmental dyslexia (DD) is a neurodevelopmental learning disorder with high heritability. A number of candidate susceptibility genes have been identified, some of which are linked to the function of the cilium, an organelle regulating left-right asymmetry development in the embryo. Furthermore, it has been suggested that disrupted left-right asymmetry of the brain may play a role in neurodevelopmental disorders such as DD.

**Methods:** Here, we studied two individuals with co-occurring *situs inversus* (SI) and DD using whole genome sequencing to identify single nucleotide variants or copy number variations of importance for DD and SI.

**Results:** Individual 1 had primary ciliary dyskinesia (PCD), a rare, autosomal recessive disorder with oto-sino-pulmonary phenotype and SI. We identified two rare nonsynonymous variants in the dynein axonemal heavy chain 5 gene (*DNAH5)*: c.7502G>C;p.(R2501P), a previously reported variant predicted to be damaging and c.12043T>G;p.(Y4015D), a novel variant predicted to be damaging. Ultrastructural analysis of the cilia revealed a lack of outer dynein arms and normal inner dynein arms. MRI of the brain revealed no significant abnormalities. Individual 2 had non-syndromic SI and DD. In individual 2, one rare variant (c.9110A>G;p.(H3037R)) in the dynein axonemal heavy chain 11 gene (*DNAH11)*, coding for another component of the outer dynein arm, was identified.

**Conclusions:** We identified the likely genetic cause of SI and PCD in one individual, and a possibly significant heterozygosity in the other, both involving dynein genes. Given the present evidence, it is unclear if the identified variants also predispose to DD, but further studies into the association are warranted.

## BACKGROUND

Left-right asymmetry conditions are characterized by failure of organization of the internal organs along the left-right axis (1). Laterality is established through a process involving motile and primary cilia at the embryonic node. A number of genes causing laterality disorders when disrupted have been identified in humans (1). About 20-25% of *situs inversus* (SI) *totalis* - a complete reversal of internal organs – individuals are also affected by primary ciliary dyskinesia (PCD) (1). PCD (OMIM #244400) is a rare autosomal recessive disorder, caused by functional impairment of the motile cilia. PCD leads to oto-sino-pulmonary disease with the phenotypic triad chronic sinusitis, bronchiectasis and SI (Kartagener syndrome) in approximately 50% of cases (2). PCD has heterogeneous underlying genetics and to date, mutations in more than 30 genes have been identified as causative, of which the dynein axonemal heavy chain 5 gene *DNAH5* accounts for the largest proportion of cases (28%) (2, 3).

Developmental dyslexia (DD) is one of the most common neurodevelopmental disorders, affecting around 5-12% of the population, and is highly heritable (4). The underlying neurodevelopmental causes of DD are not yet fully understood. One hypothesis is that neuronal migration disturbances during development lead to misplacement of neurons in the adult brain, resulting in changes in white and grey matter (4). Early studies have suggested a role in brain asymmetry, for example of the planum temporale (5).

Genetic studies of DD have led to the identification of a number of dyslexia susceptibility genes, reviewed in (6). Interestingly, some of them, namely *DYX1C1 (DNAAF4), DCDC2* and *KIAA0319*, have a reported role in cilia (7-13). In addition, loss-of-function mutations in *DYX1C1* and *DCDC2* have been found in patients showing typical ciliary deficits: *DYX1C1* in patients with PCD (14) and *DCDC2* in patients with nephronophthisis-related ciliopathy, inherited deafness and neonatal sclerosing cholangitis (15-18). Other dyslexia candidate genes, such as *CEP63* and *PCNT* are involved in centrosome and basal body biology (19, 20).

While DD has been associated with various anatomical and functional changes in the brain (21), a handful of reports have investigated brain anatomy and functionality in individuals with *situs inversus* (for example (22-26)). Interestingly, a range of neurodevelopmental disorders, such as autism, schizophrenia and DD, has been associated with laterality defects in the brain (27). The recent discoveries about ciliary genes and DD give new perspectives on the brain asymmetry theory and neuronal ciliary signaling theory in DD (13). It is currently unknown whether ciliary phenotypes and DD share a common genetic cause.

Here, we sought to address a potential common genetic cause underlying SI and DD. We studied two individuals with SI and/or PCD and DD, using whole genome sequencing (WGS) to determine a possible genetic cause for their phenotype. In addition, we performed brain imaging on one of the individuals to determine potential alterations in the brain associated with SI or DD.

## METHODS

### Sample collection and DNA extraction

DNA was extracted from saliva samples using standard procedures. For details, see Supplementary Methods.

### Whole-genome sequencing and sequence analysis

WGS was carried out at the national genomics infrastructure (NGI) at Science for Life Laboratory (SciLifeLab), Stockholm. The sequencing libraries were prepared using 1.1 µg of high quality genomic DNA samples with the Illumina (San Diego, CA) TruSeq PCR-free kits (350 bp insert size) and sequenced on one Illumina HiSeqX lane each. The data were processed using the Piper pipeline and the sequence reads were aligned to the human genome build GRCh37. Next, duplicates were removed using Picard tools and the data were recalibrated using GATK v3.3. Single nucleotide variants (SNVs) and insertions/deletions (INDELs) were called using the GATK tool HaplotypeCaller. The genetic variants were annotated using SNPEff and ANNOVAR. We were mainly interested in looking at rare homozygous or compound heterozygous loss-of-function variants in coding regions or splice variants. For downstream analysis, we extracted all the coding variants and splice variants, excluded all synonymous variants and considered further only insertions, deletions, stop-gain, stop-loss, non-synonymous and variants. The list was filtered to retain only variants with a minor allele frequency (MAF) of <1% in the 1000G (all), 1000G (European), ExAC (total) and ExAC (non-Finnish European) databases (Fig. S1, Table S1, Table S2). The impact of variants was evaluated using the prediction tools SIFT, Polyphen2, MutationTaster, CADD and GERP++. Selected variants were examined manually in the BAM files using Integrated Genomics Viewer (28). For splice effect prediction, we used SeattleSeqAnnotation138 annotation and AlamutVisual 2.11 splicing prediction module. Copy number variants (CNVs) were called using the CNVnator v.0.3.2 and both CNVs and balanced structural variants (SV) were identified using TIDDIT 2.2.3. For CNVnator the bin size was set to 1000 bp. For TIDDIT the minimum number of supporting pairs was set to 7. The human genome build GRCh37 was used for both CNVnator and TIDDIT analyses. The resulting vcf files were merged using SVDB 1.1.2. We filtered the variants by frequency using the SweGen database and included all SVs at a frequency of ≤0.001 (Table S3). Aligned sequences (BAM files) overlapping with CNVs of interest were manually inspected using the Integrated Genomics Viewer. The remaining SNVs and SVs were compared to candidate gene lists (Table S4, Table S5). For details and references to the used tools, see Supplementary Methods.

### PCR and Sanger sequencing validation

Sanger sequencing of PCR products was used to validate selected WGS variants. For details, see Supplementary Methods and Table S6.

### Electron microscopy

Electron microscopy analysis of epithelial cells from the nasal cavity was carried out using standard procedures. For details, see Supplementary Methods.

### Magnetic resonance imaging (MRI)

Individual 1 and a healthy age-matched female control (59 years old, normal-reader, right-handed) were studied. Standard clinical imaging sequences were acquired, including 3D T1- and T2-weighted images with 1.0 mm isotropic voxel size for radiological readings. We also conducted task-based functional MRI (fMRI) and diffusion tensor imaging (DTI) measurements. Details on MRI acquisition parameters and post-processing are provided in Table S7 and Supplementary Methods.

## RESULTS

### Individual 1

Individual 1 is a Swedish woman affected by PCD and DD, born to non-consanguineous parents. She presented with bronchiectasis, *situs inversus* (Fig. 1A) and has experienced recurring upper and lower airway infections since birth. Transmission electron microscopy of a nasal epithelial brush biopsy showed significant goblet cell hyperplasia, a reduced number of ciliated cells, an increased number of microvilli and sporadically distributed lymphocytes (Fig. 1B). High magnification analysis of cilia revealed a lack of outer dynein arms (ODA) (mean 2.4 +/- 0.2 per cilium; normal interval: 7-9 ODA/cilium). The number of inner dynein arms (IDA) was within normal limits (mean 4.05 +/- 0.2; normal interval: 2-7) (Fig. 1C). In addition, the individual had been diagnosed with left convex scoliosis (Fig. 1A), attention deficit hyperactivity disorder (ADHD) and Asperger’s syndrome. Furthermore, she is left-handed (Edinburgh inventory laterality index −60.00), and has the ability for mirror writing, a phenomenon overrepresented in dyslexics and left-handed persons (29). Neurological examination was normal. No history of PCD was present in the family (Fig. 1D). The father had self-reported DD and one niece was formally diagnosed with DD (Fig. 1D). For an overview of the clinical phenotype, see Table 1.

#### Whole genome sequencing

We performed WGS at an average sequencing depth of 32x. The number of total and mapped reads and the number of called variants are given in Table S1. Due to the phenotypic triad of PCD, DD and scoliosis present in the individual, we first focused our analysis on *DYX1C1*, which has previously been associated to DD, causes PCD when mutated, and absence of the *dyx1c1* orthologue causes spine curves in zebrafish (14, 30, 31). We did not find any SNVs that were previously associated to PCD or DD nor any other rare coding or noncoding variants in *DYX1C1*.

Next, we extracted the coding and canonical splice variants and filtered as described in Methods and Table S1. We focused on a set of PCD genes known to cause laterality defects (n=29) (2, 32) (S4). Due to the autosomal recessive inheritance pattern of PCD we assumed homozygosity or compound heterozygosity and looked for genes with biallelic variants. We identified two nonsynonymous variants in *DNAH5*, which were confirmed by Sanger sequencing (Fig. 1 E). The variant c.7502G>C;p.(R2501P) (rs78853309; NC_000005.9:g.13810275C>G; NM_001369.2:c.7502G>C) in exon 45 has a frequency of 4×10^−4^ in ExAC, 1.8×10^−4^ in GnomAD and has not been reported in the 1000G and SweGen databases. It is predicted to be highly conserved (GERP = 5.31) or damaging in all assessed prediction tools including a CADD score of 26.8 (see Methods and Table S2). The same variant has previously been reported in a compound heterozygote patient with PCD (3). The second variant, c.12043T>G;p.(Y4015D) (rs754466516; NC_000005.9:g.13721345T>G; NM_001369.2:c.12043T>G) in exon 71, has a frequency of 1.65×10^−5^ in ExAC, 0.81×10^−5^ in GnomAD and has not been reported in the 1000G and SweGen databases. It is conserved (GERP = 5.4) and is predicted to be damaging by all the assessed prediction tools including a CADD score of 26.5 (see Methods and Table S2). It has not previously been linked to PCD. Both of the reported variants have not been observed in a homozygous state in GnomAD. As we did not have access to parental DNA, we genotyped these variants in an unaffected sibling. The unaffected sibling is wildtype for c.12043T>G and heterozygous for c.7502G>C (Fig. 1 D), suggesting that the affected individual is compound heterozygous for the *DNAH5* variants. To test whether *DNAH5* rare variants co-segregate with DD, we performed targeted Sanger sequencing of the two nieces of the individual, one affected and one unaffected with DD. None of them carry any of the two variants in *DNAH5* (Fig. 1 D).

Upon examination of the known dyslexia susceptibility variants, we only found variants that are common in the general population (Table S5). In addition, an unbiased approach filtering by CADD score and retaining all variants with a score >25 did not reveal any additional plausible variants (Table S2). CNVs have been reported in PCD patients and in heterotaxia (33, 34) and as potential risk factors for DD (35, 36). Structural variant analysis of the WGS data revealed one inversion encompassing *CCDC103*, a gene implicated in PCD (Table S3). We did not find any other structural variants in regions overlapping with candidate genes.

#### Brain imaging

We performed MRI brain scanning to explore neuroanatomy and functionality in the presence of *situs inversus* and DD. Radiological assessment did not reveal any structural anatomical abnormalities in individual 1. The 3D T1-weighted images of individual 1 and a healthy control are presented in Fig. S2. We used fMRI in combination with a silent word generation task (Fig. 2A). We observed a bilateral activation of frontal gyri in the control, which was absent in individual 1. Interestingly, there was additional activation of the parietal lobe in individual 1. Activation clusters are detailed in Table S8. Overall, these data do not allow conclusions about the hemispheric laterality of the individual.

Next, diffusion tensor imaging (DTI) tractography of the corticospinal tracts was compared to an age- and gender-matched right-handed control subject. In individual 1, we observed more DTI tracts on the right side than on the left side, which is reversed in relation to the control. In addition, we observed fewer corticospinal tracts crossing over the corpus callosum in individual 1 compared to the control (Fig. 2 C and D). Overall, the DTI data suggest a right-dominant hemisphere in the individual, possibly linked to the left-handedness.

In summary, we identified a previously known and a novel variant in *DNAH5*, as a likely cause for PCD in this individual.

### Individual 2

Individual 2 is a Caucasian American boy affected by non-syndromic SI and DD (Gray Oral Reading Test 5 (GORT-5) overall reading index =84). He has a mild curvature of the lower thoracic spine towards the left, but no symptoms of PCD and a biopsy of the cilia was inconclusive (data not shown). His father has self-reported DD, but was not formally diagnosed (Fig. 2 A). The mother has mild scoliosis. There was no family history of SI or PCD. The individual also shows symptoms of ADHD. For an overview of the individual’s phenotypes, see Table 1.

#### Whole genome sequencing

We performed WGS at a mean sequencing depth of 25x. The total and mapped number of reads and the number of called variants are listed in Table S1.

First, we examined the *DYX1C1* gene and found the common haplotype −3G>A/1249G>T (rs3743205; rs57809907) previously associated to dyslexia (30). However, the haplotype did not co-segregate with the dyslexia phenotype, as it was also present in the mother and sibling, as revealed by Sanger sequencing (data not shown). We did not find any other SNVs that were previously associated to PCD or DD nor any other rare variants in *DYX1C1*.

As in individual 1, we extracted and filtered for coding and splice variants (see Methods and Table S1 for details on filtering) in a list of genes associated with left-right defects (Table S4). Individual 2 was heterozygous for a single rare variant c.9110A>G;p.(H3037R) (rs192327380; NC_000007.13:g.21813391A>G; NM_001277115.2:c.9110A>G) in *DNAH11* in exon 56, which we confirmed by Sanger sequencing (Fig. 3 B). The frequency of the variant is 1.5×10^−3^ in ExAc, 0.8×10^−3^ in 1000Genomes and 7.8×10^−4^ in GnomAD. Targeted Sanger sequencing revealed that this variant was inherited from the mother (Fig. 3). The variant was predicted to be benign by all of the assessed prediction tools including a CADD score of 8.4 (Table S2). We then expanded our search to more common SNVs in *DNAH11* (<50% MAF) but found none that were inherited from the father. We further checked a list of intronic variants close to exons (≤20bp), but did not find any rare variants. In addition, we looked for predicted splice effects of rare *DNAH11* deep intronic variants. We found a possible creation of a new acceptor site at chr7:21809845 (NC_000007.13:g.21809845T>G; NM_001277115.1:c.9103-3539T>G; heterozygous variant), however this variant is common (GnomAD= 6.1×10^−2^) and probably not contributing to the phenotype.

As individual 2 has non-syndromic SI, we expanded our search and looked for variants in genes known to cause L/R asymmetry defects (n=40) (1) (Table S4). We could not find any plausible disease-causing variants in these genes. There has been a report on digenic inheritance of PCD with *situs solitus* via *DNAH11* and *DNAH2* (37), so we sought for variants in *DNAH2*. We found one rare, nonsynonymous, missense variant in *DNAH2* c.2495A>G:p.(Q832R) (rs181090270; NC_000017.10:g.7646805A>G; NM_001303270:c.2495A>G; Table S2) with a frequency of 1.6×10^−3^ in the 1000Genome database, 1.5×10^−3^ in GnomAD and non-existent in ExAC, which is predicted to be benign with a CADD score of 7.167.

When examining the known dyslexia susceptibility variants we only found variants that are common in the general population (Table S5). An unbiased approach filtering all the variants with a CADD score >25 did not reveal any additional interesting variants supporting the phenotypes of individual 2 (Table S2). Structural variant analysis revealed one CNV within *NEGR1* (Table S3), a gene in which a CNV previously has been associated with DD (35). We did not find any other structural variants in regions overlapping with candidate genes.

In summary, in individual 2 we identified a common DD-associated haplotype and a rare variant of unknown significance in the outer dynein arm component *DNAH11*, both inherited from the mother.

## DISCUSSION

Several reports have demonstrated that DD candidate genes have a role in cilia (7, 8, 11, 12, 14-18). Furthermore, L/R asymmetry defects in the brain have been proposed as an anatomical basis to neurodevelopmental disorders such as schizophrenia and specifically to DD, possibly mediated by ciliary dysfunction (13, 27). Here, we studied the underlying genetic causes in two individuals with DD and SI and/or PCD. This is, to our knowledge, the first study specifically addressing the genetics of DD co-occurring with a known ciliopathy.

In individual 1, a patient with PCD and DD, we found one novel (c.12043T>G;p.(Y4015D)) and one previously reported rare variant (c.7502G>C; p.(R2501P)) in *DNAH5*, probably compound heterozygous. The *DNAH5* gene encodes one of the outer arm axonemal dynein heavy chain proteins (3). Mutations in *DNAH5* cause ODA defects, but do not affect IDAs, which is consistent with the ciliary ultrastructure found in the individual. Prediction scores support the pathogenicity of the identified variants. Expression of *DNAH5* in the developing human brain is rather low as reported in the Allen Brain Atlas and they have no known function in primary cilia/ neuronal cilia (http://www.brainspan.org/) (38). Possibly, *DNAH5* may lead to abnormal asymmetry in the brain via the left-right patterning via the cilia in the embryonic node. The asymmetry in the brain may then in turn contribute to DD. However, the genetic co-segregation pattern in the individual 1 pedigree shows that DD does not co-segregate with the variants in *DNAH5*. The DD inheritance pattern might be explained by an autosomal dominant inheritance pattern of a rare variant in another gene with incomplete penetrance in the sibling of the individual or by a complex inheritance pattern with several low penetrance common variants. WGS of all family members might clarify the inheritance pattern of DD, although the small size of the family complicates candidate-free approach WGS analysis.

The DTI tractography results suggest an inversion of hemispheric dominance, while fMRI language dominance testing was inconclusive. Abnormal symmetry of the brain - both increased asymmetry as well as decreased asymmetry – has been reported in dyslexics (5, 13, 39). Whereas some studies report a typical left hemispheric language lateralization in the SI brain, others report a reversal of the language center to the opposite hemisphere (22-26). While there is a robust association between handedness and hemispheric language dominance, the relationship between visceral situs and hemispheric dominance is more complex (25, 40, 41). It has even been suggested that brain asymmetry develops independently from the main symmetry-breaking pathway in the body (25, 41). We therefore have reason to speculate that the observed reversion of corticospinal fiber tracts is related to the left-handedness of the individual.

In summary, we identified a previously known and a novel variant in *DNAH5*, as a likely cause for PCD in individual 1.

In individual 2, we identified a haplotype previously associated to DD. However, the haplotype did not co-segregate with the dyslexia phenotype, as it was also present in the mother and sibling. After careful examination of PCD and L/R-asymmetry genes, one rare variant in *DNAH11* was identified. A causative variant in the gene *DNAH11* is consistent with the normal ultrastructure of cilia in PCD patients with *DNAH11* mutations and the clinical report that the biopsy of cilia remained inconclusive. The lack of identification of the second variant does not exclude *DNAH11* as a candidate as there might be another yet unidentified genetic variant. However, the variant is predicted to be benign and *DNAH11* mutations have not been found in 13 cases of isolated *situs inversus* without PCD (42). Furthermore, the variant c.9110A>G in *DNAH11* and c.2495A>G in *DNAH2* have been observed in a homozygous state in one individual in GnomAD. Overall, this evidence weakens the strength of *DNAH11* as a candidate in this individual. Our CNV analysis revealed no strong candidates with known links to SI or DD. In addition, it should be noted that polygenic inheritance and also environmental factors have been suggested to play a role in laterality disorders which might explain the difficulty to find the causative variants in individual 2 (1).

Taken together, we report a known and a novel variant in *DNAH5* as likely causative genetic variations for PCD that will be of value in the practice of diagnosing PCD. We believe that the reported variants are adding to the characterization of pathogenicity of SNVs. However, their involvement in DD pathology remains elusive. Regarding the DD phenotype, the possible role of these variants cannot be excluded but remains to be determined. Future functional studies should test specifically if the variants have functional consequences and aim at elucidating the role of ciliary genes on the brain in general. We propose the careful examination of variants in dynein/ciliary genes in individuals recruited for studies of DD.

## LIST OF ABBREVIATIONS

ADHD: Attention deficit hyperactivity disorder
CNV: Copy number variation
DCDC2: doublecortin domain containing 2
DD: developmental dyslexia
DNAH11: Dynein axonemal heavy chain 11
DNAH5: Dynein axonemal heavy chain 5
DTI: Diffusion tensor imaging
DYX1C1/DNAAF4: dyslexia susceptibility 1 candidate 1; dynein axonemal assembly factor 4
fMRI: Functional magnetic resonance imaging
LR: left-right
MRI: Magnetic resonance imaging
PCD: Primary ciliary dyskinesia
SI: situs inversus
SNV: single nucleotide variation
SV: Structural variant
WGS: Whole genome sequencing

## DECLARATIONS

### Ethics approval and consent to participate

Ethical permits were granted for the project by the Stockholm regional ethics committee and the ethics committee of the Central Finland Health Care District (“DYS-SWE” Ref.Nr. 2013/214-31/3; “DYSFAM” Ref.Nr. 4U/2016; Ref.Nr. 2013/1325-31/2; Ref.Nr. 2016/1684-31/1; Ref.Nr. 2016/2538-32). Written informed consent of the participants or their parents has been obtained.

### Consent for publication

Informed consent for publication was obtained by the participants or their parents.

### Availability of data and material

The datasets generated and analyzed during the current study are withheld for confidentiality reasons, but can be made available to qualified researchers by reasonable request to the corresponding authors.

### Authors’ contributions

ITP, JK and AB conceived the project. AB, EE and ITP designed and analysed sequencing experiments. AB carried out wet-lab experiments. AD performed electron microscopy and analysed data. MP performed clinical examination. HEN helped with electron microscopy analysis and provided expertise to the study. TG and TQL performed MRI acquisition and analysis. HM and JE performed CNV analysis. HM did splice effect analyses. AB wrote the manuscript. JK and AL provided expertise to the study. All authors revised, read and approved the manuscript.

## Acknowledgements

We would like to thank the participants and their family members involved in the study. We thank Ingegerd Fransson, Eira Leinonen and Auli Saarinen for help with sample handling. The authors would like to acknowledge support from Science for Life Laboratory, the National Genomics Infrastructure (NGI, Stockholm) and Uppmax, for providing assistance in massive parallel sequencing and computational infrastructure (project b2014023).

## FIGURES AND TABLES

**Table 1: Clinical, behavioral and histological characterization of the individuals**

*ODA, outer dynein arms; IDA, inner dynein arms; ADHD, attention deficit hyperactivity disorder*

**Figure 1: Phenotype and genetics individual 1**. A) X-Ray image showing *situs inversus* and left convex scoliosis. B) Low magnification electron micrograph of biopsy from nasal respiratory mucosa showing hyperplasia of goblet cells (GC), reduced number of cilia and increased number of microvilli (mv). C) High magnification electron micrograph of a cilium from the epithelial cells showing lack of outer dynein arms and normal inner dynein arms (arrowheads), normal radial spokes and central pair. Scale bar= 200 nm. D) Pedigree. Individual 1 (arrow) has PCD (black) and DD (gray). The father and the niece are affected by DD (gray). Unaffected individuals are shown in white. The genotypes of the two variants in *DNAH5* are indicated in the affected individual and in the unaffected brother (G>C denominates c.7502G>C, T>G denominates c.12043T>G, Wt denominates wildtype). E) Sanger DNA sequencing chromatogram of individual 1 and controls. F) Schematic representation of the domains of DNAH5 and localization of the amino acid substitutions p.(R2501P) and p.(Y4015D). N=N-terminus, C=C-terminus, MTB=microtubule-binding domain, P1-P6=P-loops 1-6.

**Figure 2: MRI Brain imaging individual 1**. fMRI activations overlaid on the semi-inflated FreeSurfer cortical surface for individual 1 (A) and control (B) in response to the silent word generation task. DTI tractography of corticospinal tracts of individual 1 (C) and the matched control (D). Right hemisphere tracts in blue, left hemisphere tracts in green. Volume of interest in red.

**Figure 3: Genetics individual 2**. A) Pedigree. Individual 2 (arrow) has *situs inversus* (black) and DD (gray). The father has self-reported DD, but the dyslexia diagnosis is not confirmed (question mark). Unaffected individuals are shown in white. The genotypes of the variant in *DNAH11* are indicated (A>G denominates c.9110A>G, Wt denominates wildtype). B) Sanger DNA sequencing chromatogram of individual 2 and controls (2 unaffected siblings). C) Schematic representation of the domain structure of DNAH11 and localization of the amino acid substitution p.(H3037R). N=N-terminus, C=C-terminus, P1-P4= P-loops 1-4, MTB= microtubule domain, AAA1-6= AAA modules 1-6.

