## Supplementary Material for "Rare variants in dynein heavy chain genes in two individuals with *situs inversus* and developmental dyslexia"

*Contents:*


- Supplementary Methods
- Figure S1. WGS analysis workflow
- Figure S2. Structural MRI findings.


*Attachments:*


- Table S1. Filtering strategy
- Table S2. SNVs: Short listed variants
  - 2A) Candidate genes
  - 2B) Unbiased approach via CADD filtering
- Table S3. SVs: Structural variants
  - 3A) Short list
  - 3B) Full list
- Table S4. PCD and L/R genes used for filtering
- Table S5. Dyslexia variants
  - 5A) List of variants associated with dyslexia, reading, spelling or other literacy-related language traits
  - 5B) Short list of dyslexia-associated exonic variants
- Table S6. List of PCR Primer sequences
- Table S7. MRI acquisition parameters
- Table S8. fMRI activation clusters in silent word generation task

### SUPPLEMENTARY METHODS

#### Electron microscopy

Epithelial cells from the nasal cavity were collected by gentle brushing and immediately fixed in 2.5% glutaraldehyde in 0.1 M sodium cacodylate buffer. The cells were collected by centrifugation and the resulting pellet was further fixed in osmium tetroxide, dehydrated in a graded ethanol series, and embedded in epoxy resin. Ultrathin sections were cut and contrasted with uranyl acetate and lead citrate. Digital images were captured with a FEI Tecnai BioTwin transmission electron microscope (FEI Inc., Eindhoven, Netherlands) equipped with a Gatan Orius SC1000 CCD camera (Gatan Inc., Abingdon, United Kingdom). Data are expressed as mean +/- SEM.

#### Sample collection and DNA extraction

Saliva samples were collected using Oragene DNA kits (OG-500; DNA Genotek, Ottawa, Canada) and genomic DNA from saliva was extracted using the prepIT PT-L2P (DNA Genotek) according to the manufacturer´s instructions. DNA concentration and purity were measured using Nanodrop 8000 (Thermo Fisher Scientific, Waltham, MA) and Quant-iT dsDNA BR assay kit with a Qubit fluorometer (Thermo Fisher Scientific). High molecular weight of the DNA was verified on an agarose gel.

#### Whole-genome sequencing and sequence analysis

The data were processed using the Piper pipeline (www.github.com/NationalGenomicsInfrastructure/piper). The sequence reads were aligned to the human genome build 37 using bwa_mem v0.7.12 (https://github.com/lh3/bwa) (https://arxiv.org/abs/1303.3997) Duplicates were removed using Picard tools (http://broadinstitute.github.io/picard/) and the data were recalibrated using GATK v3.3 (https://software.broadinstitute.org/gatk/) (1). Single nucleotide variants (SNVs) and insertions/deletions (INDELs) were called using the GATK tool HaplotypeCaller. The genetic variants were annotated using SNPEff and ANNOVAR through http://wannovar.wglab.org/ (2, 3). For CNV and SV calling, we used CNVnator v.0.3.2 (4) and TIDDIT 2.2.3 (5). The databases used for filtering were 1000 genomes project (6), available at http://www.internationalgenome.org, accessed June/September 2017 and Exome aggregation consortium (7), available at http://exac.broadinstitute.org, accessed June/September 2017. We consulted GnomAD database, available at http://gnomad.broadinstitute.org/. For individual 1, additionally, the SweGen database was consulted (8) available at https://swefreq.nbis.se, accessed July 2018. For splice effect prediction, we used SeattleSeqAnnotation138 annotation (http://snp.gs.washington.edu/SeattleSeqAnnotation138/)(9) and AlamutVisual 2.11 splicing prediction module (https://www.interactive-biosoftware.com/alamut-visual/) (10). The impact of variants was evaluated using the prediction tools SIFT (11), Polyphen2 (12), MutationTaster (13), CADD (14) and GERP++ (15).

**PCR and Sanger sequencing validation**

Sanger sequencing of PCR products was used to validate selected WGS variants. PCR assays were run according to standard protocols using HotStarTaq Plus DNA Polymerase (203605, Qiagen, Hilden, Germany). The products were purified from 1% agarose gels using QIAquick Spin PCR purification kit (28106, Qiagen) or Nucleospin Gel and PCR Clean-up kit (740609, Macherey-Nagel, Duren, Germany) according to the manufacturer´s instructions. The Sanger sequencing was performed at Eurofins Genomics (Ebersdorf, Germany). The list of primers used for the PCRs is provided in Table S6.

**Assessment of handedness**

Individual 1 was self-reported ambidextrous with a tendency to left-handedness. She was therefore assessed further by the adapted Edinburgh handedness inventory (16) using the online tool http://www.brainmapping.org/shared/Edinburgh.php. Individual 2 and the healthy control in the MRI experiment were self-reported right-handed.

**Magnetic resonance imaging (MRI)**

All MRI scans were conducted on a whole-body 3T Prisma$^{Fit}$ clinical MRI scanner (Siemens, Erlangen, Germany) using a 64-channel head coil. Both the fMRI and DTI scanning protocols were adapted to the acquisition protocols in the Human Connectome Project with 2 mm isotropic voxel size (www.humanconnectomeproject.org/). Acquisition parameters are detailed in Table S7.

The task-based fMRI consisted of three runs of word generation conducted using a block design. The task was 4 epochs of silent word generation interleaved with resting periods. Each task epoch was 32 s. The task cue was either standard English alphabet letters or non-letter unreadable symbols displayed in front of the subject with an MRI-compatible screen. In response to a letter, the participant was instructed to generate as many words as possible starting with that letter in silence. In response to an unreadable symbol, the participant was instructed to rest.

The DTI session included 3 runs (each lasting approximately 7 minutes), representing 3 different gradient tables. Each gradient table includes approximately 90 diffusion weighting directions plus 6 b=0 acquisitions interspersed throughout each run. Diffusion weighting consisted of 3 shells of b=1000, 2000, and 3000 s/mm$^2$ interspersed with an approximately equal number of acquisitions on each shell within each run. The gradient tables were identical to those used for the Human Connectome Project (www.humanconnectomeproject.org/).

**fMRI post-processing**

The fMRI datasets underwent a pre-processing procedure, which was performed with the **A**nalysis of **F**unctional **N**euro**I**mages (AFNI) (Version Debian-16.2.07~dfsg.1-3~nd14.04+1, http://afni.nimh.nih.gov/afni) and FSL (FIMRIB Software Library) (http://www.fmrib.ox.ac.uk/fsl) programs with a bash wrapper shell. After temporal de-spiking, six-parameter rigid body image registration was performed for motion correction. The average volume for each motion-corrected time series was used to generate a brain mask to minimize the inclusion of extra-cerebral tissues. Spatial normalization to the standard MNI template was performed using a 12-parameter affine transformation and mutual-information cost function. Nuisance signal removal was performed by voxel-wise regression using 14 regressors based on the motion correction parameters, the average signal of the ventricles and their 1$^{st}$ order derivatives. After baseline, trend removal up to the third order polynomial effective band-pass filtering was performed using low-pass filtering at 0.08 Hz. Local Gaussian smoothing up to FWHM = 3mm was performed using an eroded gray matter mask. For the task-based fMRI datasets, we performed generalized linear model (GLM) type of regression analysis using the stimulus timing parameters and derived linear regression coefficients for the stimuli. To assess the difference in fMRI activation between the control and the affected individual we conducted ANOVA analyses of the GLM regression results for the 3 repeated sessions using the AFNI program 3dANOVA. The statistical significance of the activations was assessed by using a voxel-wise threshold of $p<0.001$ and a cluster size > 100 spatially connected voxels.

**DTI post-processing**

The diffusion weighted images were eddy-current corrected using the average T2-weighted spin echo images as reference with FSL program eddy-correct. After rigid motion correction, the DTI data were spatially normalized to MNI template using affine registration described above for the fMRI data. The diffusion tensor elements were then estimated using the AFNI program 3dDWItoDT. With the tensor elements we conducted whole-brain white matter stratigraphy using the program dtiquery (http://graphics.stanford.edu/projects/dti/software). To determine the laterality of the main motor tracts, volumes of interest were placed in each of the corticospinal tracts on the right and left side.

### SUPPLEMENTARY FIGURES

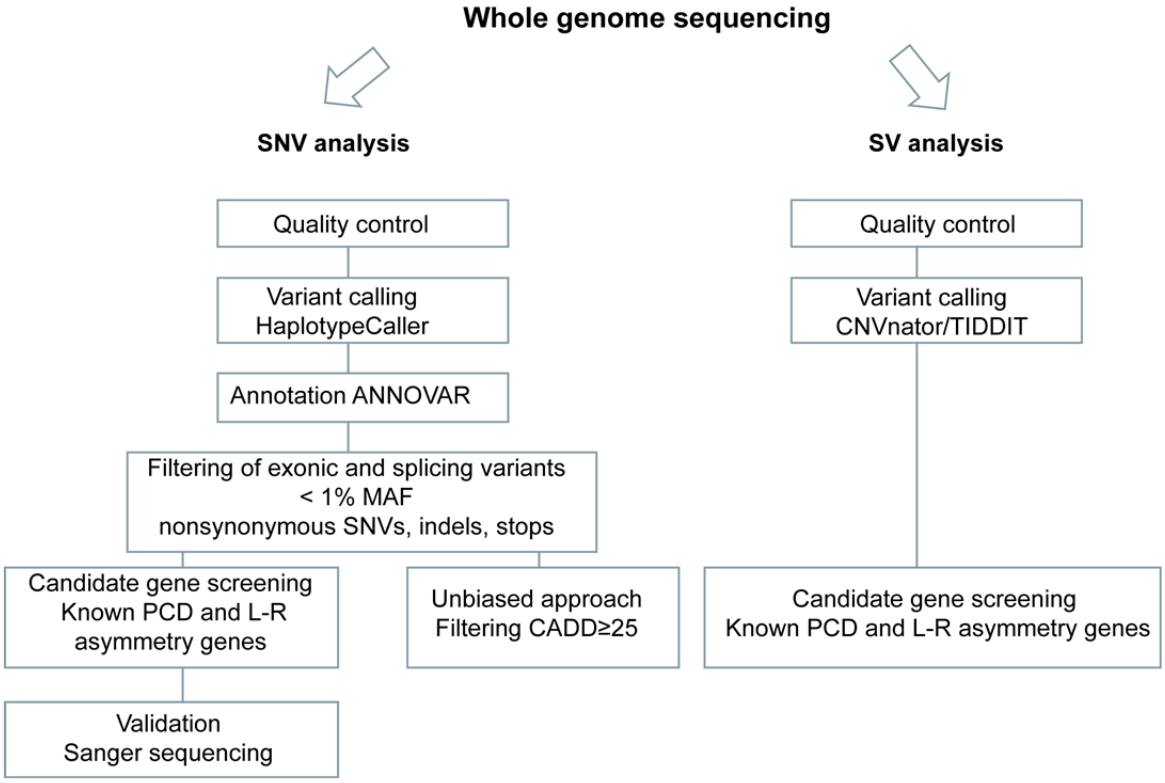


**Fig. S1: WGS analysis workflow**

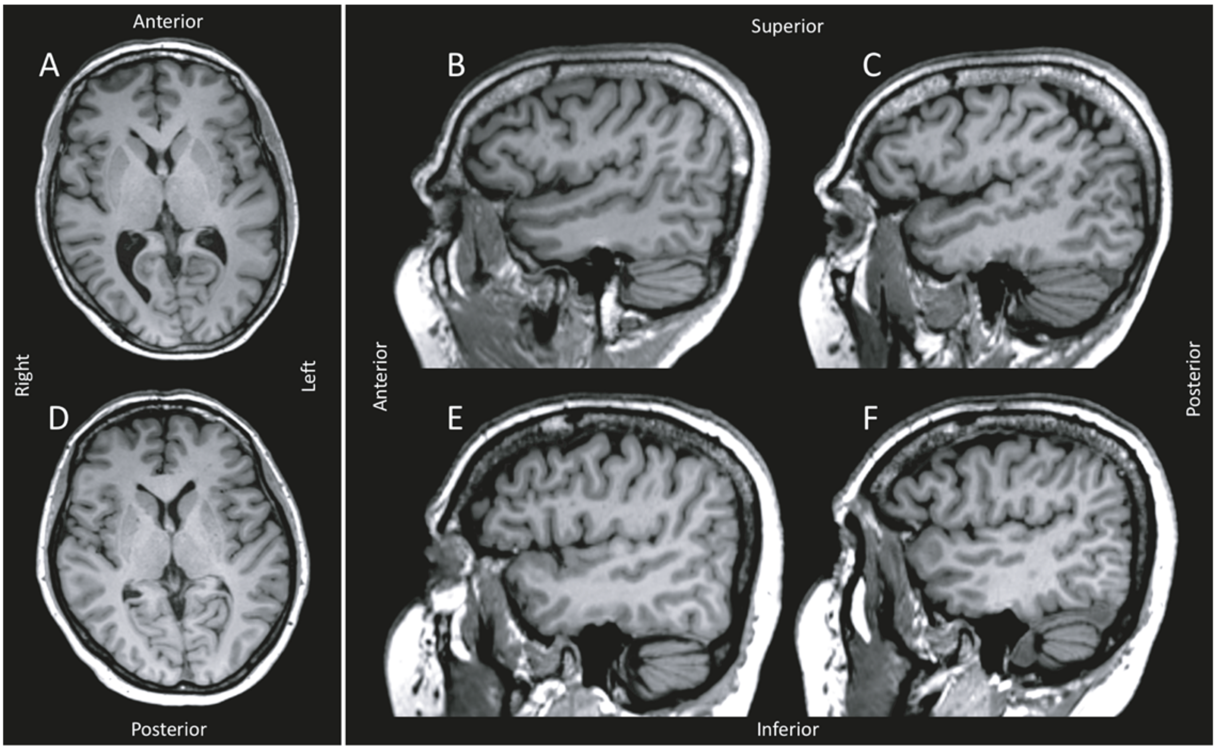


**Fig. S2: Structural MRI findings.** Axial (first column) and sagittal (second and third column) 3D T1-weighted images of proband 1 (top row) and the matched healthy control (bottom row). Radiological readings did not reveal any significant structural anatomical anomalies.
